## Supplementary Information for "Surface remodeling and inversion of cell-matrix interactions underlies community recognition and dispersal in *Vibrio cholerae* biofilms"

**This PDF file includes:**
Supplementary Methods
Supplementary Figures S1 to S15
Supplementary Tables S1 to S3

### Supplementary Methods

### Molecular dynamics simulations

We utilize coarse-grained molecular dynamics (CGMD) simulations to investigate the interaction between polymers and bacterial cells, focusing on its impact on biofilm structure. We employ the OpenFSI package by Ye and colleagues[1], designed for efficient fluid-structure interaction (FSI) simulations and implemented by Large-scale Atomic/Molecular Massively Parallel Simulator (LAMMPS)[2]. The bacterial cell is simulated with a lattice membrane model with a dimension of 3 by 1 μm (length by diameter). We discretize the membrane (the bacterial surface) into a triangular network of 1179 vertices and 2354 elements, maintaining an approximately uniform Lagrangian mesh with an average size of 90 nanometers. To represent the polymers in the system, coarse-grained (CG) beads with a diameter of 100 nm are introduced to the simulation box. The mechanical characteristics of the membrane are defined by applying potential functions to the membrane's triangular networks. These functions' parameters are based on our recent work[3]. Specifics regarding these potential functions are described in Ye *et al.*,[1]. For interparticle interactions, we use Lennard-Jones (LJ) potential with various interaction modes among the beads and nodes on the bacterial surface. The interaction details are summarized in the Supplementary Table S2.

The radial distribution function (RDF) quantifies the number density in the radial direction ($r$) between nodes from different cells. This calculation involves dividing the radial channel into uniform bins of size d$r$. Due to the anisotropic nature of the cell geometry, we calculate the node–node distance between different cells to determine the cell–cell distance instead of using the center-of-mass distance. We count the number of nodes locating inside these bins and denote it as $n_{\mathrm{NP}}$. Finally, $g(r) = \frac{n_{\mathrm{NP}}}{V \times N_{\mathrm{nodes}}}$ is computed, where $V$ represents the bin volume and $N_{\mathrm{nodes}}$ represents the total number of nodes in the channel. The appearance of the peak at 1.14 μm in the RDF profile indicates a strong parallel alignment of the rod-shaped bacterial cells. This feature served as a criterion to distinguish between a depletion-aggregated and a well-dispersed system. When the cells are bridged by polymers, the sharp peak at 1.14 μm in the RDF profile is replaced by a broader peak with the peak position shifted to a higher value. This occurs because bridging increases the distance between cells, resulting in a higher cell-to-cell separation.

The bacterial cells are randomly positioned within a simulation box of 6.2 × 6.2 × 6.2 μm$^3$, in which periodic boundary conditions are applied in the $x$, $y$, and $z$ directions.

To systematically study the depletion behavior of bacterial cells (Fig. S7a), we vary three parameters: the number of cells, the number of polymer beads, and VPS surface coverage. VPS surface coverage is adjusted by randomly replacing native cell surface beads with VPS-coated surface beads. The interaction between VPS beads and cellular surface nodes depends on the type of node: native cell surface nodes exhibit repulsive interaction with the polymer beads (1 $k_{\mathrm{B}}T$), while VPS-coated surface nodes have a much weaker repulsion (0.1 $k_{\mathrm{B}}T$). The number of cells ranges from 6 to 45, corresponding to volume fractions from 5.4% to 39.4%. The number of VPS polymer beads varies between 30,000 and 100,000, representing volume fractions of 6.6% to 21.9%. The percentage of VPS-coated beads ranges from 0% (no VPS coating) to 100% (full VPS coating).

To investigate the impact of VPS bead properties on the depletion process (Fig. S7b-d), we use a model of 12 bacterial cells interacting with 80,000 VPS beads. We switch the interaction between the cell surface beads and the VPS beads from repulsion to strong attraction with an attraction strength of 50 $k_{\mathrm{B}}T$. Additionally, the interaction between the VPS beads is also changed from pure repulsion to strong attraction, allowing us to observe the effect of VPS bead properties (attractive or repulsive) on cell aggregation. To incorporate the attractive interactions, we extend the cut-off distance for the LJ potentials to 2.5 in the simulations. We also vary different attraction strengths (18 $k_{\mathrm{B}}T$, 30 $k_{\mathrm{B}}T$, and 50 $k_{\mathrm{B}}T$) to study the effect of attraction strengths on cell bridging (Fig. S7e). To examine the effect of VPS polymer bead concentrations on bridging aggregation, we reduce the number of attractive VPS beads from 80,000 to 25,000 (Fig. S7f). Finally, to achieve a transition between depletion and bridging, we randomly select a portion of the VPS beads (up to 20%, or 16,000 beads) in the system containing 12 cells and 80,000 VPS beads and convert them to 'sticky' beads. These sticky beads can attract both the cell surface beads and each other while remaining repulsive to the native VPS beads. The disappearance of the sharp peak at 1.14 μm in the RDF profiles, which signifies the depletion of bacterial cells, is used to track the transition between depletion and bridging (Fig. 4d).

In all simulations, we employ the Langevin thermostat to model the interaction of bacterial cells and polymers with a background implicit solvent. This thermostat is combined with the NVE (conserving number of atoms – volume –

total energy) ensemble to perform Brownian dynamics with the damping factor of 100. The temperature of the system is maintained at 300 K in all models. To ensure the phase separation is completed, a long simulation for each model is performed with a maximum of 100,000,000 running steps. The timestep used in all the models is set at 0.01 (11.4 ns). As a result, the maximum runtime for a depletion model is calculated at 100,000,000 × 11.4 ns = 1.14 s. All the general parameters of CG modeling are summarized in Table S2. All snapshots of CGMD simulations are rendered using VMD[4].

### Mathematical modeling of spatial stochastic dynamics of biofilms

*1/ Model assumptions and parameterization*:
We model the population of *V. cholerae* cells by using a square 2D grid of $N \times N$ spots. Each spot can be empty, or occupied by a bacterium, which can be a producer (P) or a non-producer cell (nP). The dynamics proceed as a sequence of updates that include cell divisions, death/wash out events, and conversion between the two types. Variables used in the model are summarized in Table S3.

*System size and cell density*: Cells are spaced roughly with a distance of 1-3 µm apart. The fact that the vertical system size (several layers of cells, or 20-30 µm) is much smaller than the horizontal dimension, motivates the 2D approximation used here, which integrates out the dynamics along the vertical axis. While computationally less expensive, 2D simulations capture the essence of the system in this context.

*Cell division*: Division happens by picking a random neighbor out of the 4 neighbors and placing a cell of the same type in that spot, unless that spot is already occupied. If it is occupied, reproduction does not happen. P cells produce a substance (not explicitly modelled) that allows them to stick to the surface and to each other. This substance is costly, meaning that the division rate P is reduced due to the fact that they use resources to produce the substance. Denote by $r_{\mathrm{p}}$ and $r_{\mathrm{np}}$ the division rates of P and nP cells, respectively. The reproductive fitness advantage of nP cells is denoted by $s$ such that $r_{\mathrm{np}} = (1+s)\, r_{\mathrm{p}}$. The measured cell replication rate of every 30 mins translates into the division rate of $r_{\mathrm{p}} = \ln 2/0.5 = 1.39\ \mathrm{hrs}^{-1}$. We further assume that the relative advantage of nP cells is $s = 0.2$ (based on prior measurement[5]).

*Cell death and washing out*: We assume that all bacteria die at a certain rate, denoted by *d*. In the absence of a flow, the two cell types have the same death rate, which is small compared to the division rate ($d = 0.05\, r_{\mathrm{p}}$). In the presence of a flow, we assume that nP cells are washed out from the system at a constant rate, which is flow-dependent. The wash-out rate of a P cell depends on how many P neighbors this P cell has. The more P neighbors that surround a P cell, the less its wash-out rate becomes. Note that P cells are assumed to have a smaller wash-out rate than nP cells even in the absence of P neighbors due to cell-surface adhesion. The total removal rate consists of the death rate *d* and a wash-out rate (in the presence of a flow):

$$d_{\mathrm{nP}} = \begin{cases} d, & \text{no fow} \\ d + d_{\mathrm{f}}, & \text{with flow} \end{cases}, \quad d_{\mathrm{P}} = \begin{cases} d, & \text{no fow} \\ d + A d_{\mathrm{f}}\left(1 - \frac{k}{\upsilon}\right), & \text{with flow} \end{cases}$$

Here *k* is the number of P cells among the neighbors of a given P cell, and coefficient $\upsilon = 5$ is used to characterize the adhesion force of P cells to each other. The constant $0 < A < 1$ indicates that even if a P cell has no P neighbors, its wash-out rate ($Ad_{\mathrm{f}}$) is still smaller than that of a nP cell: $Ad_{\mathrm{f}} < d_{\mathrm{f}}$.

*The process of conversion*: The conversion from P→nP cells and nP→P cells happens in a density-dependent manner. It is assumed that the amount of nutrients available is negatively correlated with the total number of cells. A lack of nutrients promotes the P→nP conversion, which is modeled as:

$$\mu_{\mathrm{p}\to\mathrm{nP}} = \mu \left(\frac{k'}{N^2}\right)^{\beta}$$

where $k'$ is the total number of cells (P or nP) in the system, $k'/N^2$ is their density, and $\beta = 2$. To estimate $\mu$, we assume that the maximum conversion rate is such that 90% of cells are converted within 4 hours (measured experimentally). This results in $\mu = |\ln 0.1|/4 = 0.58\ \mathrm{hrs}^{-1}$. In our simulations, we used values of $\mu$ ranging from 0.1 to 0.58 $\mathrm{hrs}^{-1}$. In contrast, the conversion nP→P increases with the amount of available nutrients:

$$\mu_{\mathrm{nP}\rightarrow\mathrm{P}} = \mu\left(1 - \frac{k'}{N^2}\right)^{\beta}$$

*2/ Stochastic simulations:*

In the images shown in this paper, the grid size is $50 \times 50$, with periodic boundary conditions. The grid is initialized by placing a P cell with probability 0.15 and a nP cell with probability 0.1. The simulations are carried out based on the stochastic simulation algorithm (SSA)[6]. More precisely, given a state of the system X, every possible event $m$ has a given propensity $a_m(\mathrm{X})$ based on the model's rates. The time at which the next reaction $m$ will occur is exponentially distributed with intensity $a_m(\mathrm{X})$. Thus if $r_m$ is a uniform random number in [0,1], setting the time for the next $m$-reaction as $\tau_m = -\log(r_m) / a_m(\mathrm{X})$ will generate random times with the correct probability distribution. This algorithm produces exact solutions to the master equation resulting from the specified rates. In particular, this means that each simulated outcome (one run of the algorithm) occurs with the correct statistical frequency.

We code the algorithm using indexed priority queues implemented with binary heaps to find the next stochastic event. We do so using the so-called Next Reaction Method[7]. We also use indexed priority queues to update the effects of nearby neighbors on a given individual. As a result, the simulations are fast and amenable to large populations, with an overall algorithmic complexity of $\mathcal{O}(n \log n)$, where $n$ is the number of individual cells.

In our simulations, we are interested in parameter sensitivities of our system as well as discovering parameter values that give a resemblance to qualitative experimental data. Parameters of key significance are $\mu$ (the maximum conversion rate) and $\beta$ (exponent in conversion rate). For example, given $\mu = 0.58\mathrm{hrs}^{-1}$, we find that $\beta = 0.1$ leads to total population collapse in the presence of flow. However, increasing $\beta$ to 2 produces nonzero steady-state populations for both P and nP cells in the presence of flow. These parameter values also produce spatial images, in steady state, that resemble experimental images of *V. cholerae* biofilm after a wash-out event. A full analysis of the parameter space will be reported in a future paper.

*3/ Producers and cheaters*:

This modeling framework allows further exploration of different assumptions. We have assumed above that nP cells do not participate in the secretion of adhesive matrix and do not benefit from it, but can be interconverted with the P cells. A different type of cells, which we term Cheaters (C, referring to the usual terminology of the evolutionary game theory), have the ability to utilize the matrix to gain benefit from its adhesive effect. In this modified version of the model, C will take on the role of nP, requiring only a change to the total removal rate in the presence of flow. The total removal rate of C cells is comprised of the death rate $d$ and wash-out rate (in the presence of a flow):

$$d_{\mathrm{C}} = \begin{cases} d, & \text{no fow} \\ d + d_{\mathrm{f}}\left(1 - \frac{k}{v}\right), & \text{with flow} \end{cases}$$

Using numerical simulations, we have explored differences between the co-dynamics of P and nP cells on the one hand, and P and C cells on the other. To do this we separately ran simulations with P and nP cells, and simulations with P and C cells. Fig. S13a plots the time-series simulations of P with nP (green and black lines) and P with C (blue and black lines). The steady states in the absence of flow are nearly identical in both cases, and we observe that P cells comprise a small minority of the population. After the flow is applied, we observe that the balance is reversed, and now P cells are the majority. The proportion of C cells in the presence of flow (under all parameters being equal) is larger compared to the proportion of nP cells (the black line is above the red line).

This trend is investigated more systematically in Fig. S13b, where we plot the steady state density of nP cells and C cells for different values of $d_{\mathrm{f}}$ (the flow). As mentioned above, the two solutions coincide in the absence of flow ($d_{\mathrm{f}} = 0$), and they also become identical for very large values of $d_{\mathrm{f}}$, where all cells are washed out. For intermediate flow values, there are more C cells, compared to the number of nP cells that could be maintained in the system. The reason for this phenomenon is their ability to utilize the matrix and stick to their P neighbors. Therefore, this simulation supports our hypothesis that the inherent cell-VPS repulsion facilitates cell dispersal by ensuring that cells that no longer secrete VPS can be removed by environmental flows.

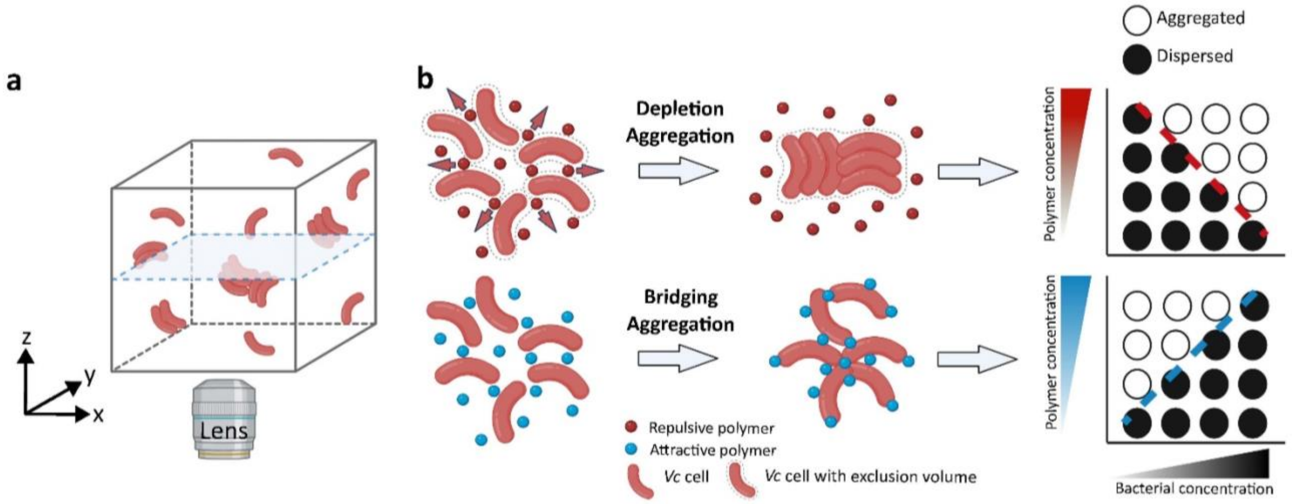

**Fig. S1 Aggregation of *V. cholerae* (*Vc*) cells in various polymer solutions.** (**a**) Schematic depicting the experimental setup for imaging nonmotile *Vc* cells (red) in a polymer solution by using a confocal microscope. (**b**) Schematic of the difference between depletion- and bridging-aggregation. *Top* (depletion-aggregation): In the presence of repulsive, non-absorbing polymers, as two cells approach one another, polymers are excluded (arrows), increasing the volume available to the polymers and the entropy of the entire system. Dashed lines represent the excluded volume around each cell. *Bottom* (bridging-aggregation): In the presence of attractive polymers, the polymer molecules can adsorb simultaneously on more than one cells and bringing them into proximity. Moreover, features on a 2D phase diagram of polymer concentration versus bacterial concentration can be used to identify the aggregation mechanism. The schematic (adapted and modified from Secor *et al.*, 2018[8]) on the right shows that bacteria remain dispersed when bacterial numbers and/or polymer concentrations are low (filled circles). Aggregation occurs when bacterial numbers and/or polymer concentrations increases (open circles). The red dashed line with a negative slope indicates the phase boundary separating the dispersed and aggregated regions in the case of depletion-aggregation, whereas the blue dashed line with a positive slope indicates the phase boundary in the case of bridging-aggregation.

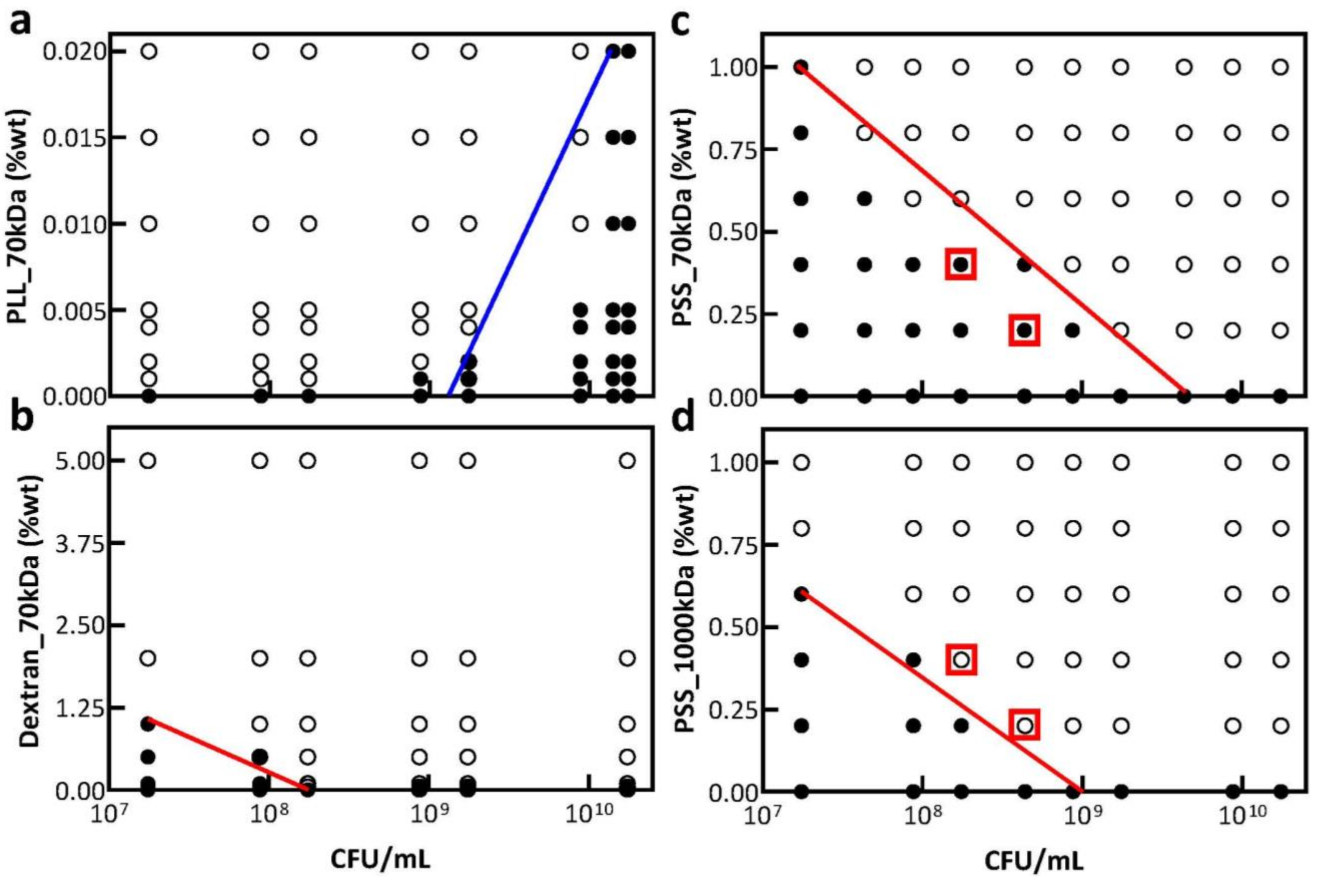

**Fig. S2 Various polymers with different molecular weights (MWs) can cause bridging- or depletion-aggregation of *Vc* cells.** Phase diagrams were generated by mixing non-VPS-producing cells (5Δ cells) and polymers at the indicated concentrations and visually scoring the cultures as either aggregated (open circles) or dispersed (filled circles) after 6 hours. Dashed lines indicate phase boundaries; a positively sloped phase boundary (blue line) indicates bridging-aggregation; a negatively sloped phase boundary (red line) indicates depletion-aggregation. Several polymers were used: (**a**) Poly-L-lysine (PLL, positively charged, MW = 70 kDa); (**b**) Dextran (uncharged, MW = 70 kDa); (**c-d**) Polystyrene sulfonate (PSS, negatively charged). Two MWs were used, 70 kDa in (c) and 1000 kDa in (d). The two red squares depict comparable bacterial and polymer concentrations in (c) and (d), with opposite phase behaviors, showing enhanced tendency of depletion-aggregation as the polymer MW increases. %wt = weight percent.

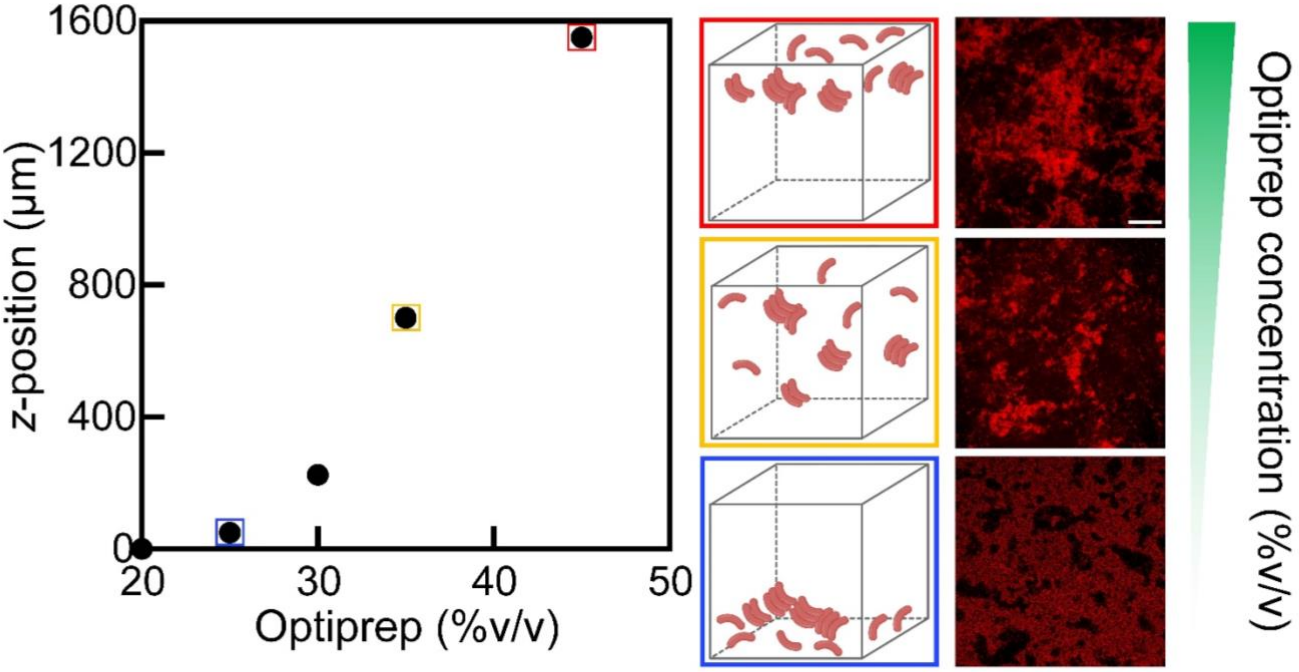

**Fig. S3 Spontaneous depletion-aggregation in VPS-producing culture is not caused by gravity.** One potential confounding factor in interpreting aggregation in VPS-producing cultures is gravity: bacterial cells are slightly heavier than the growth medium and may accumulate at the bottom of the imaging well due to gravity. Solutions with various Optiprep concentrations were therefore used to adjust the density of the LB medium to exclude sedimentation as the mechanism for aggregation. *Left*: *z*-position of the center of biomass distribution in an imaging well at different Optiprep concentrations (%v/v). *Right*: Representative images captured 24 hours after growing Δ*ABC* cells constitutively expressing mNeonGreen (pseudocolored in red) in media with various Optiprep concentrations (%v/v). The schematic for each configuration is shown on the left of the images, with the color of the schematic border corresponding to the Optiprep concentration points shown on the graph. Scale bar = 20 μm. Regardless of the density of the medium and the *z*-location of the cells, spontaneous aggregation occurs indicating that it is not caused by gravity.

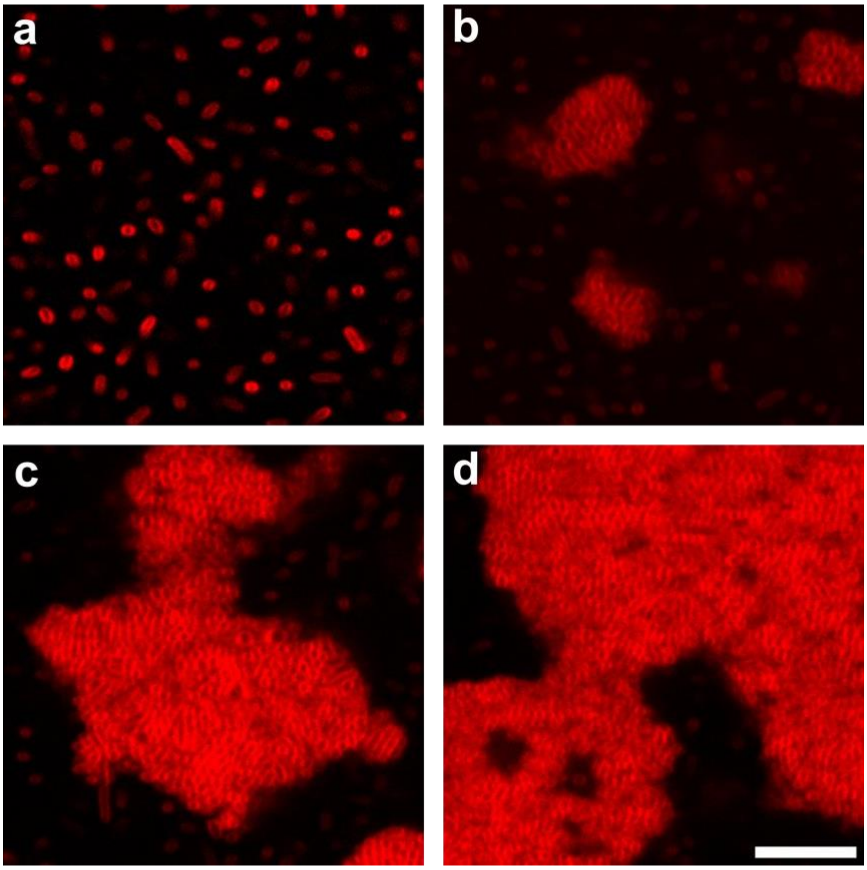

**Fig. S4 VPS is not attractive to *Vc* cells.** (**a-d**) Zoom-in cross-sectional confocal images of Fig. 1b (at $z$ = 4 μm above the glass surface) after (a) 8, (b) 12, (c) 13 and (d) 14.5 hours of biofilm growth. Cell membranes were stained with FM 4-64 (2 μg/mL). Scale bar = 10 μm.

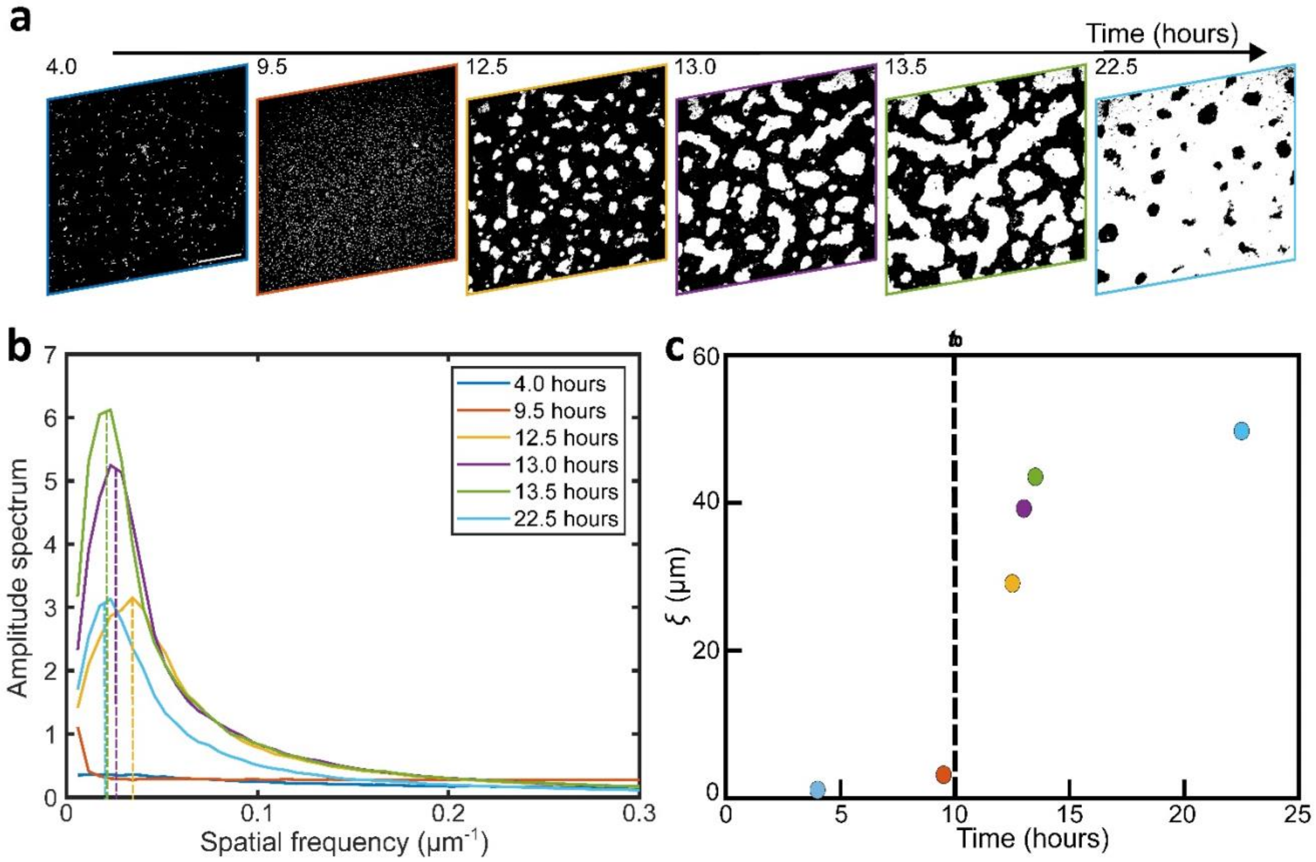

**Fig. S5 Analysis of large-scale phase separation dynamics of Δ*ABC* culture.** (**a**) Time-lapsed binarized confocal images at $z$ = 4 μm during the growth of a culture from a VPS-producing strain (Δ*ABC*). Scale bar = 50 μm. (**b**) Amplitude spectra for representative images in (a) versus spatial frequency obtained using a Fast Fourier transfer (FFT) method. The characteristic length $\xi$ was extracted from the inverse of the spatial frequency at the highest amplitude for each curve (indicated by the dashed lines). (**c**) Evolution of $\xi$ versus time $t$ in one Δ*ABC* culture. The color of the image border in (a) corresponds to the time points of the corresponding colors in (b) and (c).

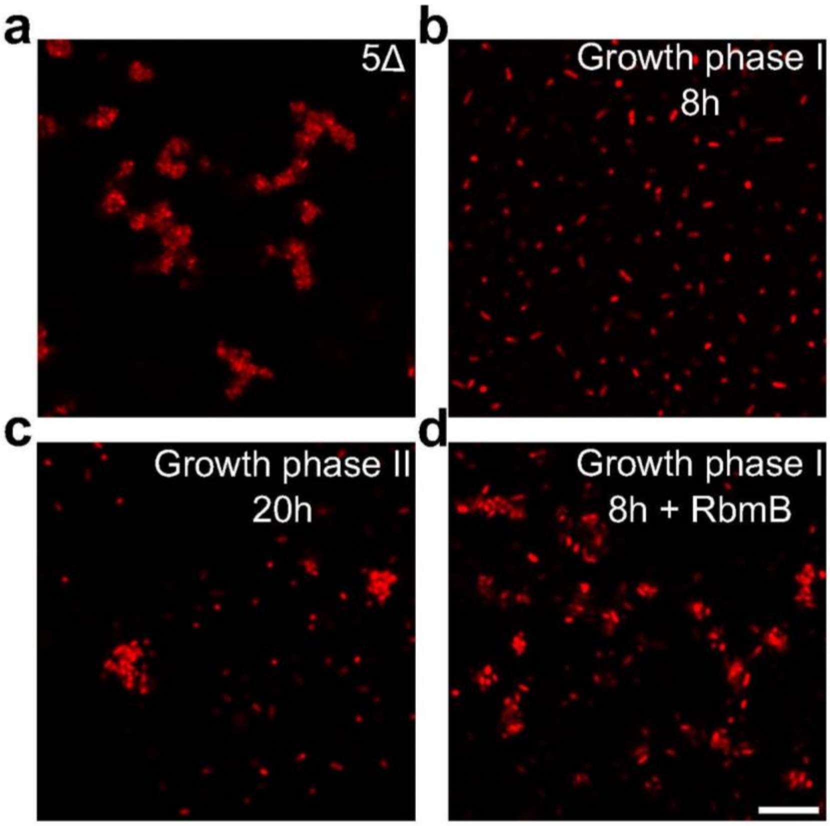

**Fig. S6 Surface remodeling underlies spontaneous depletion-aggregation.** Representative confocal images were collected after mixing purified VPS (pVPS) and (**a**) non-matrix-producing cells (5Δ cells), (**b**) matrix-producing cells (Δ*ABC* cells) chemically fixed in growth phase I (8h of growth), (**c**) chemically fixed in growth phase II (20h of growth) and (**d**) chemically fixed in growth phase I and incubated overnight with 0.05 mg/mL RbmB (RbmB was subsequently removed during the washing step). Cell concentration used in this data set equals $1.75 \times 10^8$ CFU/mL and the VPS concentration equals to (a): 0.5; (b) 0.15; (c) 0.6 and (d) 0.6 mg/mL. We observed depletion aggregation in all cases except (b). This data set is representative of raw data used to generate the phase diagram in main Fig. 2a-c. All cells constitutively express mRuby3. Scale bar = 10 µm.

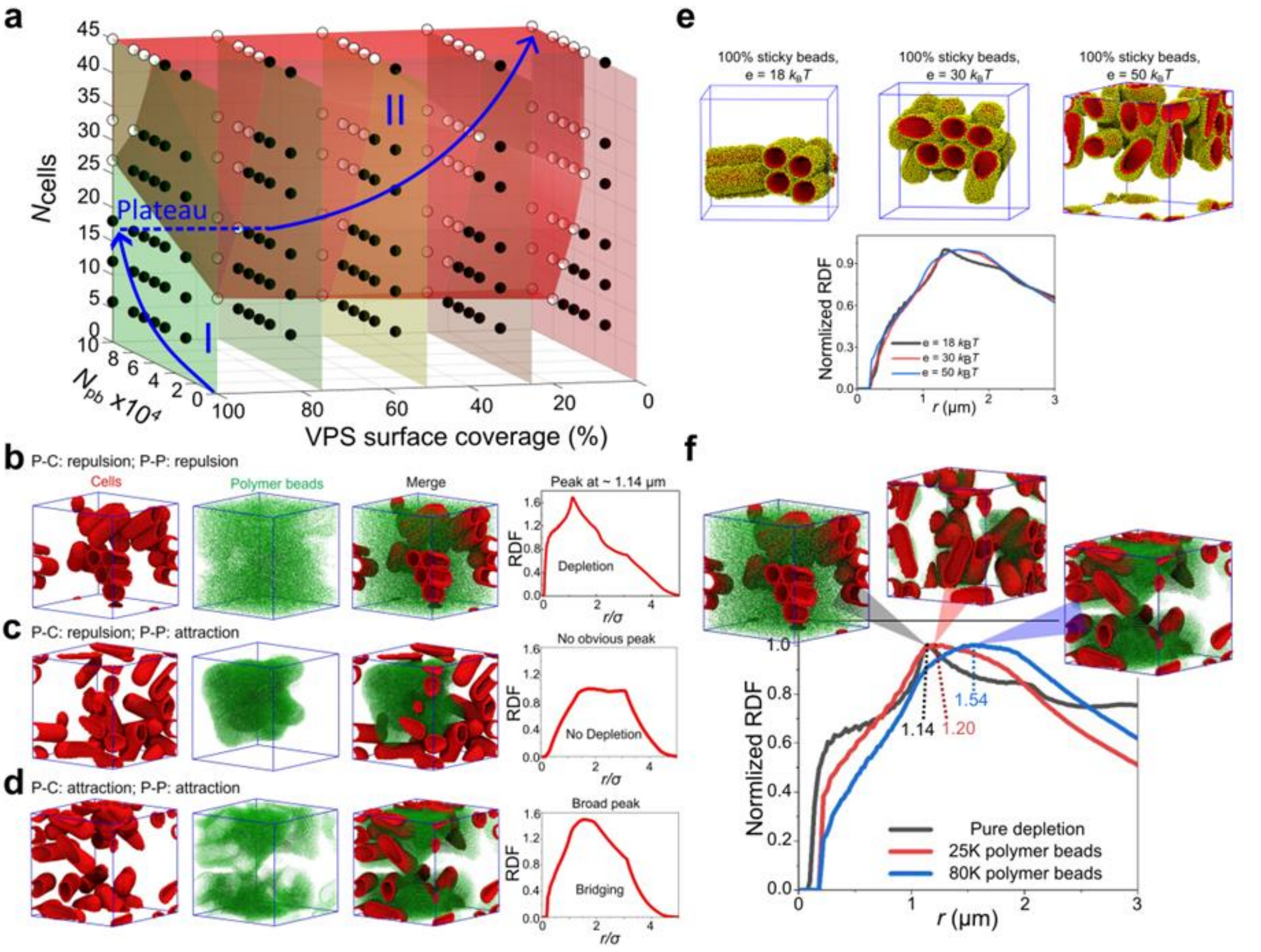

**Fig. S7 Simulations on polymer-mediated cell-cell interaction.** (**a**) A 3D phase diagram that conceptually rationalizes the developmental trajectory of the Δ*ABC* culture that produces non-crosslinked VPS. The same data was shown as in main Fig. 2g, but now in three-dimensional space. The ranges considered for the number of cells, number of polymer beads (pb), and VPS surface coverage are from 6 to 45 cells, 30,000 to 100,000 beads, and 0 to 100% VPS surface coverage, respectively. The blue line is a guide to eye for the developmental trajectory of a VPS-producing culture on the 3D phase diagram. During the 1st growth phase, both cell number and VPS concentration increase, but the system remains in the single-phase region. Upon nutrient limitation, cells are no longer coated by VPS, so the system crosses the 2D phase boundary. The increase in cell number during the 2nd growth phase further pushes the system deep into the aggregated region. (**b-d**) Coarse-grained simulation results for a system showing depletion-aggregation (b), no aggregation (c), and bridging-aggregation (d), depending on the interaction between polymers (P) and cells (C). *Left*: Snapshots of cellular (red) and polymer (green) arrangements after 100 million simulation steps. *Right*: The corresponding radial distribution function (RDF). There is an emerging peak characteristic of depletion-induced parallel arrangement of the rod-shaped cells at 1.14 μm (b), and a broad peak characteristic of bridging-aggregation (d). (**e**) Effect of attraction strength on bridging aggregation. Stronger attraction strengths lead to disordered aggregates similar to those experimentally observed. For visual clarity, only those polymer beads within a distance of 0.2 μm from the cell surfaces are shown in this panel (in yellow). (**f**) Effect of polymer bead concentrations on the bridging-aggregation. At low number of polymer beads (25,000), the RDF has a broad peak but the peak position is close to the depletion case. Simulation snapshot (*inset*) shows that one layer of polymer beads is sandwiched between and directly bridges neighboring cells. As the number of polymer beads in the simulation box increases to 80,000, the RDF becomes broader and the peak position is shifted to a larger distance. Simulation snapshot shows that clusters of polymer beads are bridging neighboring cells. For details of the simulation parameters, refer to Supplementary Table S2.

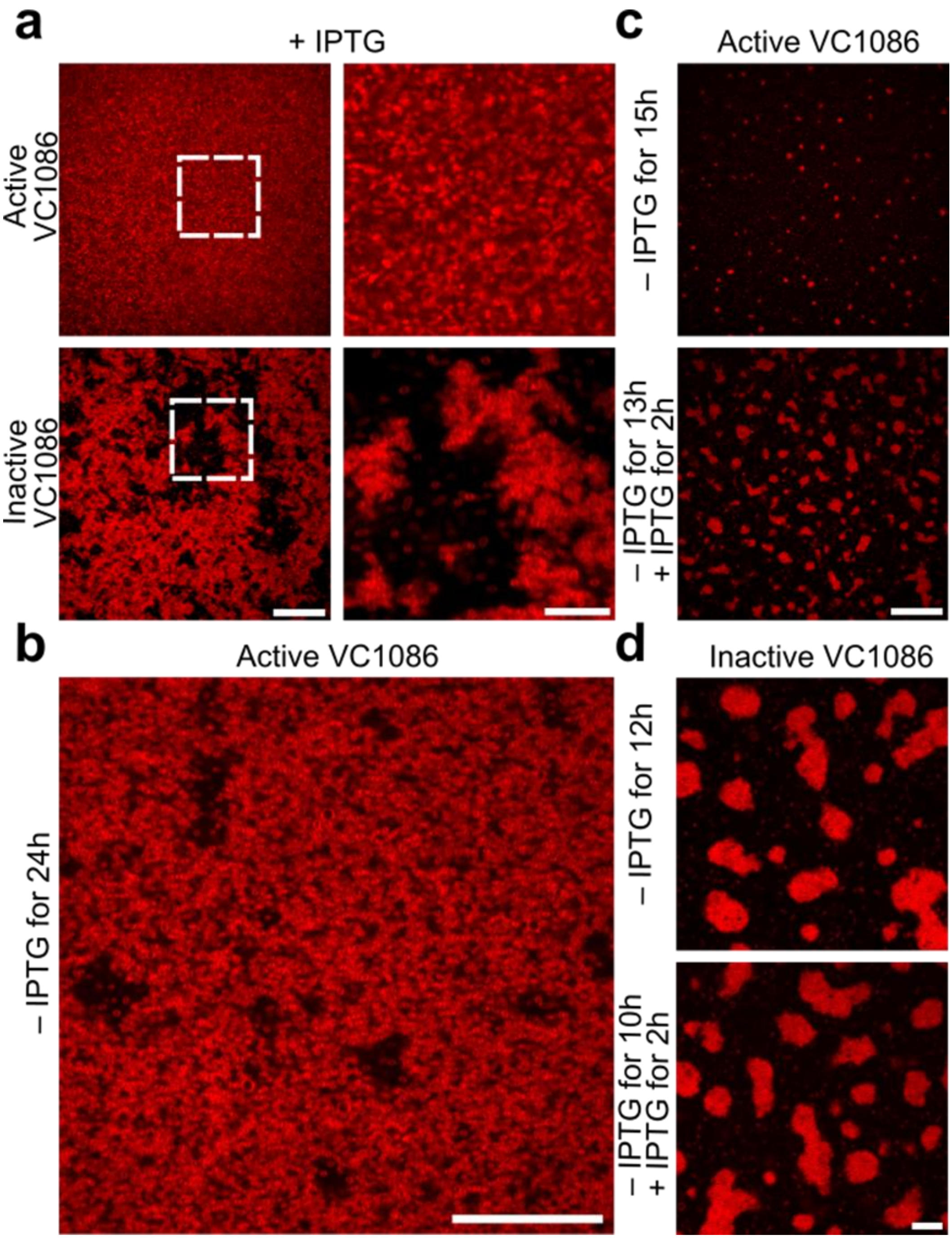

**Fig. S8 Controlled transition of depletion-aggregation.** (**a**) *Left* (the same data as main Fig. 2d): Cross-sectional confocal images of cell culture from a strain carrying a plasmid with (*top*) IPTG-inducible expression of VC1086, a c-di-GMP phosphodiesterase, and an isogenic strain carrying a control plasmid with an inactive version of VC1086 (*bottom*). Scale bar = 30 µm. *Right*: Zoom-in image of the region indicated by the white square. Both strains were grown overnight in the presence of 1 µM IPTG. Scale bar = 10 µm. (**b**) Cross-sectional confocal images of a cell culture of the strain carrying the active VC1086, after 24 hours of growth without IPTG, showing depletion aggregation. Scale bar = 15 µm. (**c**) The same data as main Fig. 2e: The strain with active VC1086 was grown for 15 hours without IPTG (*top*), or for 13 hours without IPTG followed by 2 hours of IPTG treatment (*bottom*). (**d**) The strain with inactive VC1086 was grown under the same conditions as (c). No difference was observed with or without IPTG. Note that due to the leakage of the IPTG inducible promoter in *Vc*[9,10], even without IPTG the spontaneous aggregation is delayed in the strain with active VC1086. Therefore, at 12h where the parental strain has already formed large depletion-aggregates, the strain with active VC1086 has just begun to phase separate due to the lowered level of c-di-GMP. Consequently, we had to perform this experiment at 13-15 h for (c). This was not an issue for the strain with inactive VC1086 so the experiment in (d) was performed at an earlier time according to the phase separation dynamics of the parental strain. In all experiments, cell membranes were stained with FM 4-64. Scale bar = 10 µm. All images were taken at $z$ = 4 µm above the glass surface.

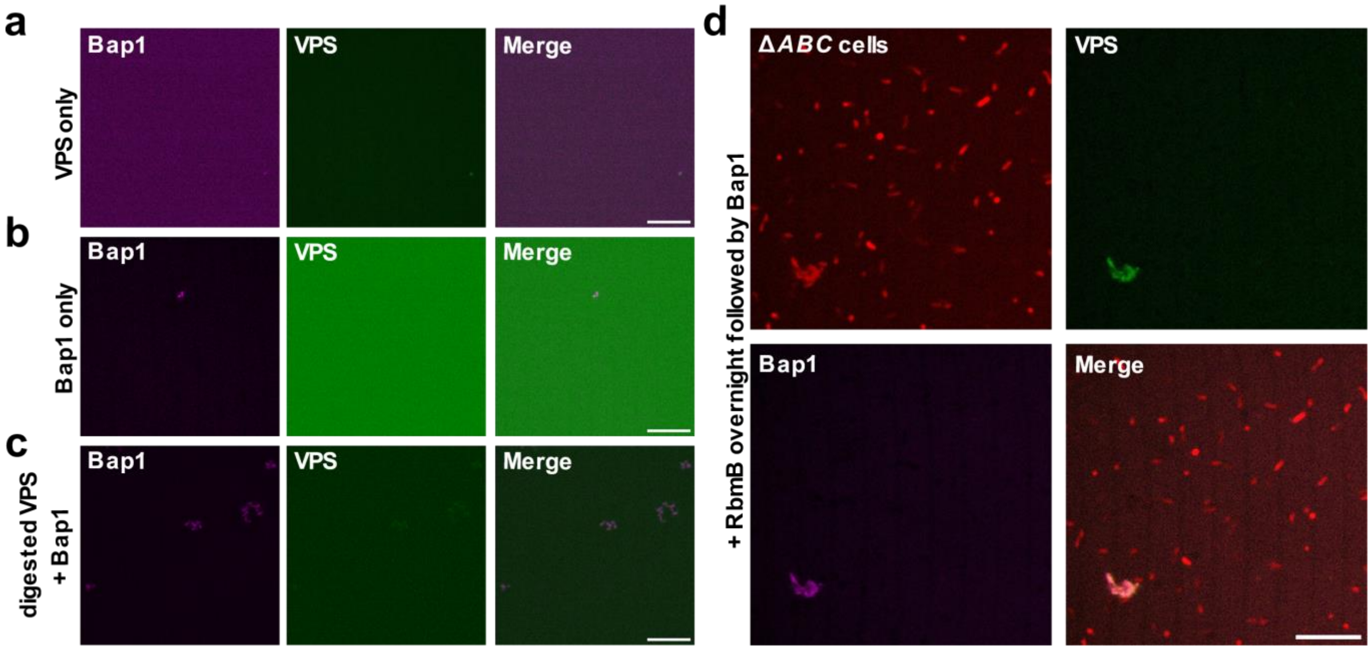

**Fig. S9 Absence of aggregation with pVPS or Bap1 alone; RbmB disrupts cell-matrix interactions.** (**a**) Cross-sectional images from a solution of pVPS at 0.6 mg/mL stained with WGA conjugated to AlexaFluor647 (pseudocolored in green). (**b**) Cross-sectional images from a solution of Bap1-GFP at 1.2 mg/mL (pseudocolored in magenta). (**c**) Cross-sectional images from a solution of pVPS at 0.6 mg/mL digested overnight with RbmB (0.05 mg/mL), washed and then mixed with Bap1-GFP at 1.2 mg/mL. Images in (a-c) were taken at $z$ = 4 μm above the glass surface); scale bars = 10 μm. No clumps were observed when either pVPS or Bap1 was present alone in solution. Additionally, after overnight digestion with RbmB, pVPS cannot form clumps when mixed with Bap1. (**d**) Cross-sectional images (at $z$ = 10 μm above the glass surface) of Δ*ABC* cells chemically fixed in growth phase I (8h after growth), mixed with RbmB overnight (0.05 mg/mL), washed and then mixed with Bap1-GFP at 1.2 mg/mL (pseudocolored in magenta). Cell membranes were stained with FM 4-64 (red) and VPS was stained with WGA-AlexaFluor647 (pseudocolored in green). Scale bars = 10 μm. The VPS signal disappears, and cell clusters are disassembled, showing that Bap1 alone cannot bridge cells if VPS molecules on the cell surfaces are trimmed off.

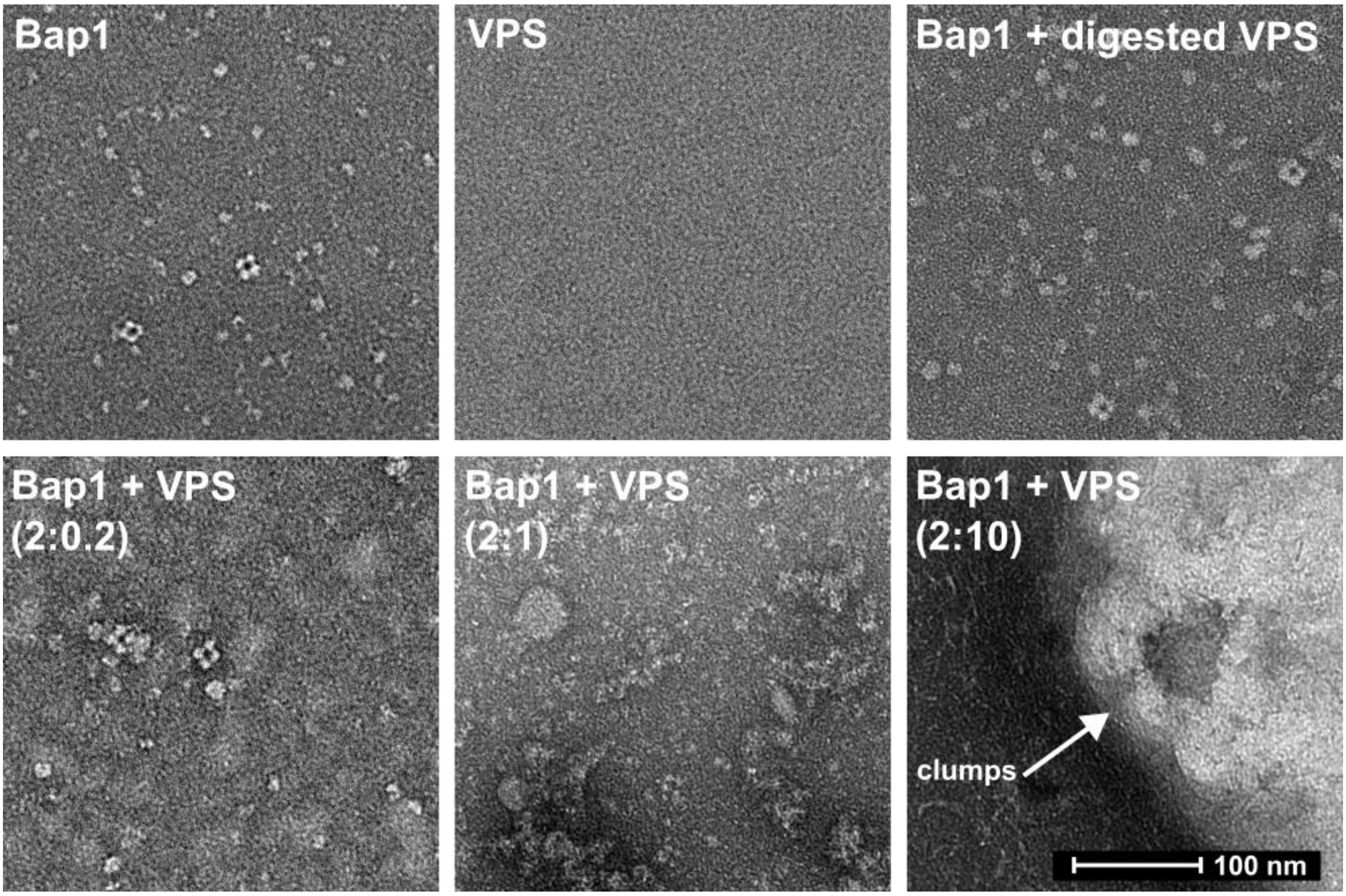

**Fig. S10 RbmB digests VPS *in vitro*.** Negative-staining electron microscopy images of Bap1 (*top left*), VPS (*top middle*) alone, and their complexes (*bottom row*). VPS does not have sufficient contrast in electron microscopy and therefore cannot be visualized. Clumps of increasing sizes are observed as VPS:Bap1 ratio increases, while keeping Bap1 concentration constant. Incubation with RbmB degrades the clumps (*top right*).

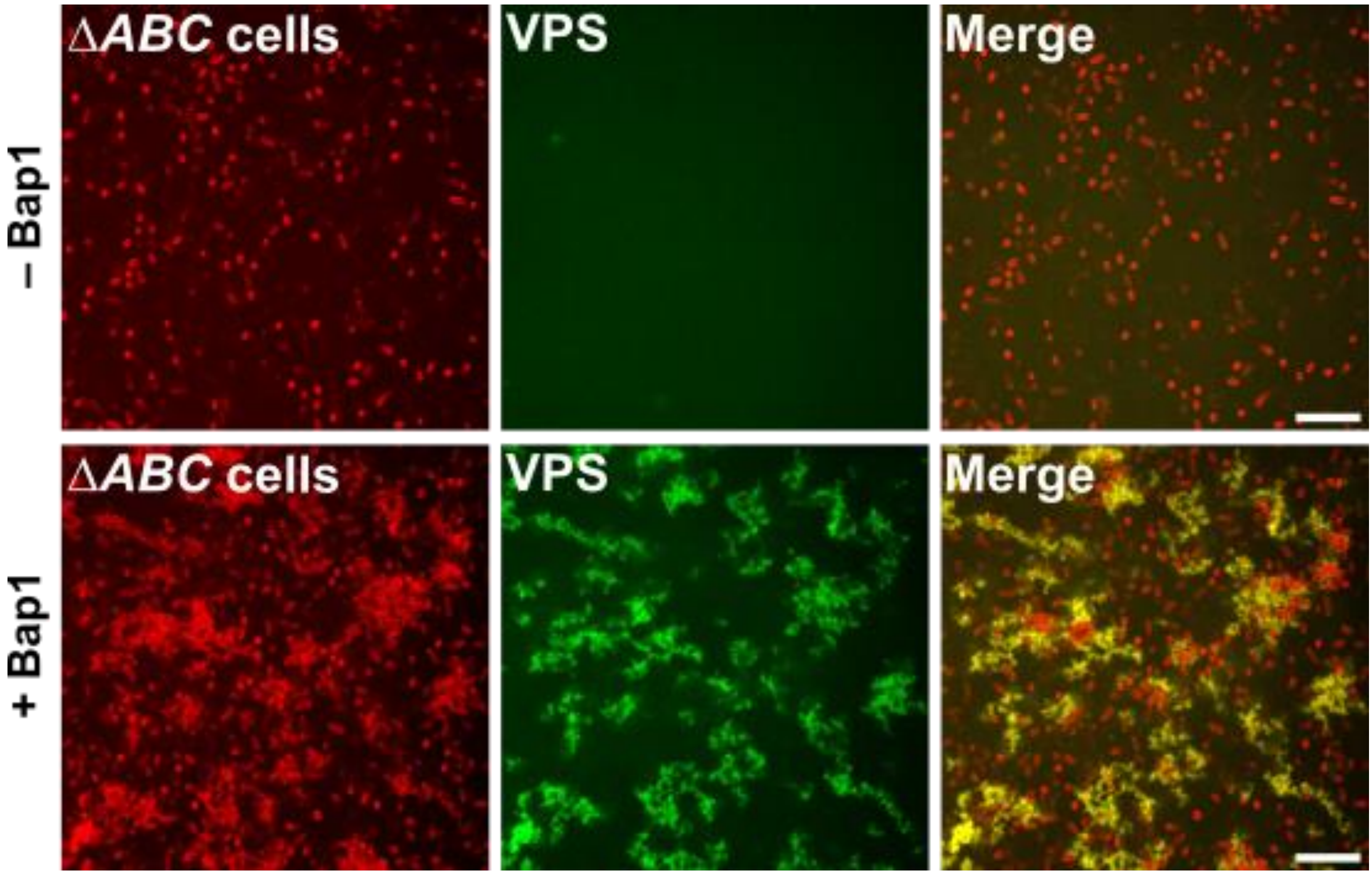

**Fig. S11 Crosslinking of VPS by Bap1 leads to bridging-aggregation.** Large view of the cross-sectional images shown in Fig. 3b of Δ*ABC* culture after 8 hours of growth, with or without 1 mg/mL Bap1 (images taken at $z$ = 4 μm above the glass surface). Cell membranes were stained with FM 4-64 (red), VPS was stained with wheat germ agglutinin (WGA) conjugated to AlexaFluor647 (pseudocolored in green). Scale bar = 10μm.

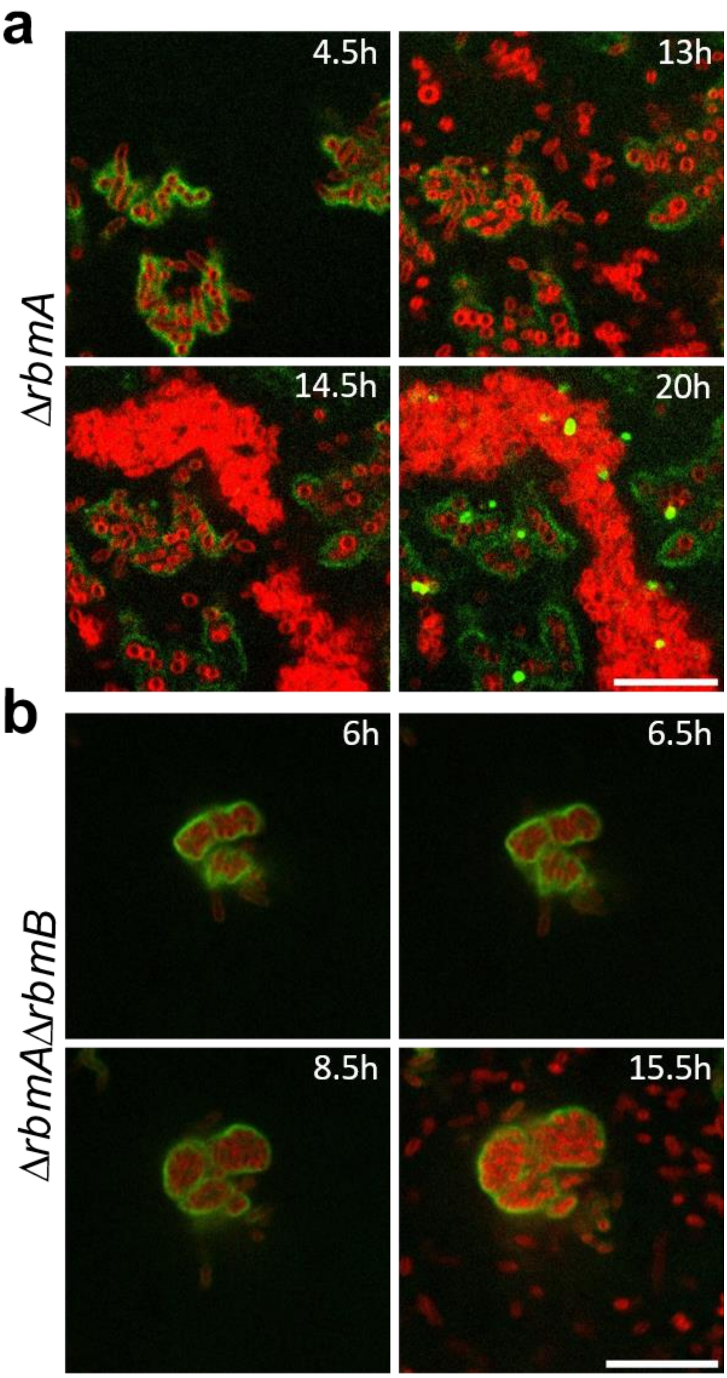

**Fig. S12 Inversion of cell-matrix interaction during biofilm growth.** (**a**) Shown are zoomed-in views of the images presented in Fig. 4a during a time-course growth from 4.5 to 20 hours of a biofilm formed by the Δ*rbmA* strain (images shown at $z$ = 4 μm above the glass surface). Cell membranes were stained with FM 4-64 (red) and VPS was stained with WGA conjugated to Oregon Green (green). Note at the late stage, some dead cells with exposed cell walls were also stained with WGA. Scale bar = 10 μm. (**b**) Zoomed-in views of the cross-sectional images presented in Fig. 5a, at $z$ = 4 μm above the glass surface from a time course of biofilm growth of Δ*rbmA*Δ*rbmB*. From 6.5 to 8.5 hours, the aggregate of cells surrounded by VPS grows in size, while dispersed cells outside of this aggregate not surrounded by VPS are visible at 15.5 hours. In contrast with the aggregates not surrounded by VPS in (a), these dispersed cells do not form any depletion aggregates. From this difference we infer that no free VPS polymer is present in the solution of Δ*rbmA*Δ*rbmB* biofilm due to the absence of the trimming effect of RbmB. Cell membranes were stained with FM 4-64 (red) and VPS was stained with WGA-Oregon Green (green). Scale bar = 10 μm.

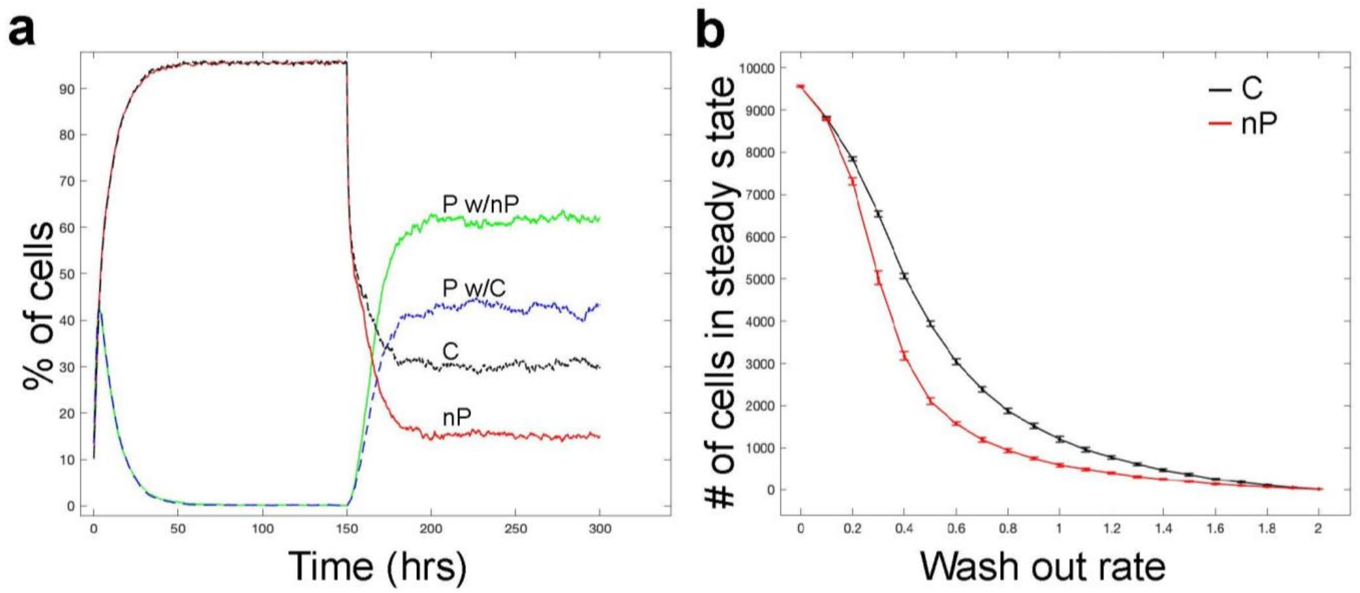

**Fig. S13 Simulated dynamics of biofilms containing producer (P) and non-producer (nP) or cheater (C) cells**. (**a**) Dual time-series plot of simulations of P with nP (green and red curves, respectively) and P with C cells (blue and black curves, respectively). This result demonstrates the advantage that cheater cells have over non-producer cells in the presence of a flow since cheater cells have a reduced wash-out rate depending on the number of surrounding P neighbors. (**b**) Plots of the average steady-state number of C and nP cells as a function of the wash out rate $d_f$. C and nP cells have, on average, the same steady state value in the absence of a flow ($d_f = 0$) and in the presence of high flow ($d_f = 2$); however, at intermediate flow rates, it is more difficult for C cells to be removed by flow. Parameter values used for (a-b): $A = 0.5, \beta = 2, \mu = 0.1, r_{\mathrm{P}} = 1.39, r_{\mathrm{nP}} = 1.2 \times 1.39, d = 0.05 r_{\mathrm{P}}, d_{\mathrm{f}} = 0.6.$

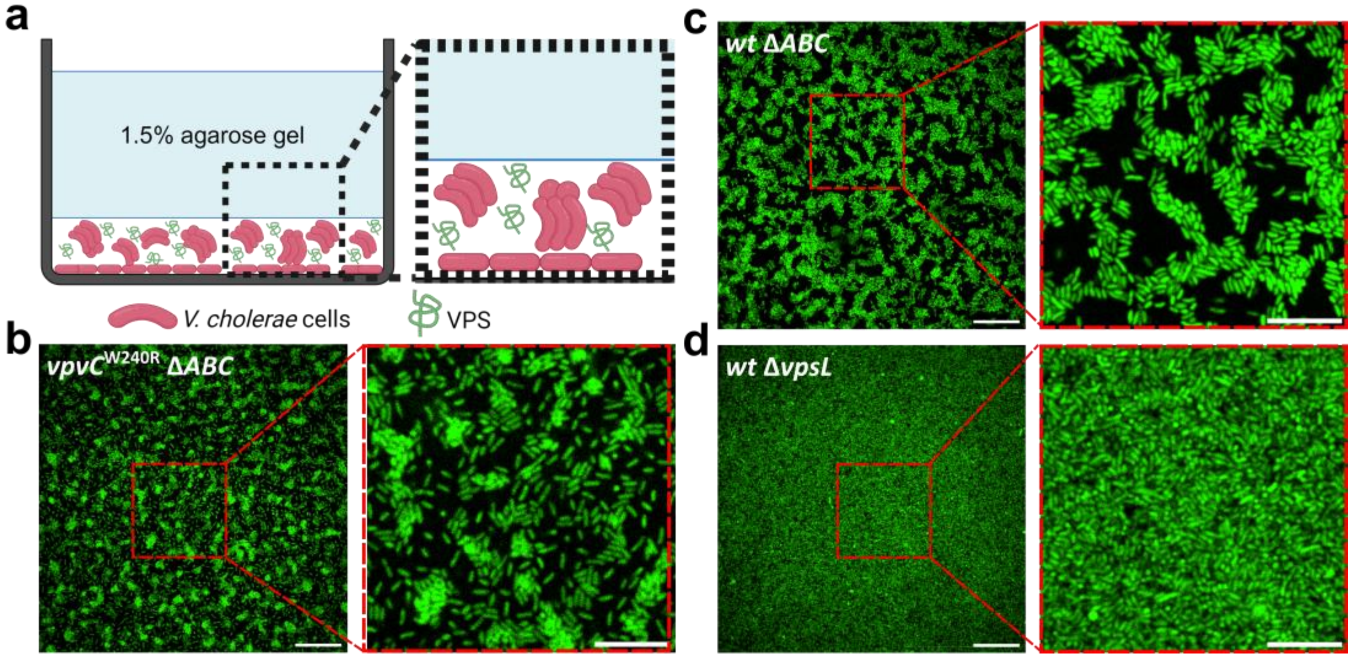

**Fig. S14 Spontaneous depletion-aggregation in the wild-type background.** (**a**) Schematic representation of cell localization and VPS molecules during biofilm development with confinement by a 1.5% agarose gel. VPS polymer is unable to diffuse through the confining agarose gel due to the small pore size. (**b-d**) Cross-sectional images ($z$ = 2 µm above the glass surface) of cultures of *vpvC*$^{W240R}$ Δ*ABC* cells (b), *wt* Δ*ABC* cells (c), and *wt* Δ*vpsL* cells (d) after 24 hours of growth in LB, all constitutively expressing mNeonGreen. Spontaneous depletion-aggregation is observed in the wild-type background under confinement of the 1.5% agarose gel, driven by an increase in polymer concentration. In contrast, cells unable to produce VPS (*wt* Δ*vpsL*) do not exhibit aggregation. Scale bars equal 50 µm on the left and 10 µm in the zoom-in images on the right.

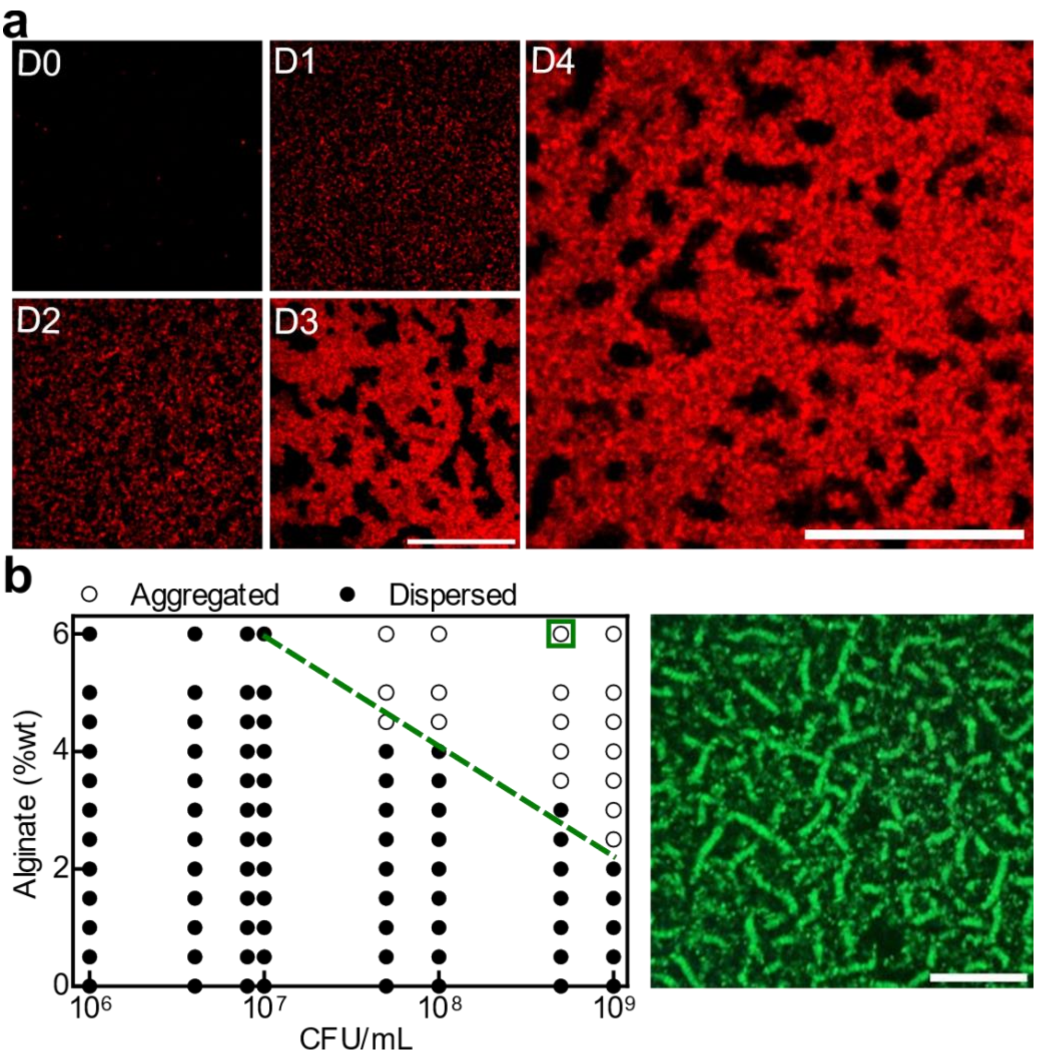

**Fig. S15 Spontaneous depletion-aggregation in mucoid *Pseudomonas aeruginosa* (*Pa*) strains.** (**a**) Cross-sectional confocal images during 5-day-growth of a culture from a mucoid strain of *Pa* (PAO1Δ*mucA*) at $z$ = 4 µm above the glass surface. Cell membranes were stained with FM 4-64. Spontaneous aggregation was observed starting from Day 2. Non-alginate producing strains such as PAO1 and PA14 did not show spontaneous depletion-aggregation (data not shown). Scale bars = 50 µm. (**b**) To test the interaction between the *Pa* cells and alginate, we used a PAO1Δ*pel*Δ*psl* strain that does not produce any EPS. *Left*: Phase diagram was generated by mixing alginate and PAO1Δ*pel*Δ*psl* cells, at the indicated concentrations and visually scoring the cultures as either aggregated (open circles) or dispersed (filled circles) after 6 hours. *Right*: Representative confocal image collected 6h after mixing alginate and PAO1Δ*pel*Δ*psl* cells constitutively expressing GFP (at $z$ = 4 µm above the glass surface). Scale bar = 20 µm. A similar phase diagram with negative sloped boundary was observed, indicating the repulsive nature between the cell and alginate. While this set of experiments indicates the possibility that similar evolution of cell-matrix interactions may happen in biofilms made by certain *Pa* strains, further research needs to be performed to test the extent to which the analogy holds and how the presence of other matrix components (such as Psl) affects the aggregation process.

**Table S1:** *V. cholerae* strains used in this study

| **Strain** | **Genotype** | **Source** |
|---|---|---|
| AM001 | *vpvC*$^{W240R}$Δ*rbmA*Δ*bap1*Δ*rbmC*Δ*vpsL*Δ*pomA*Δ*VC1807*::$P_{tac}$-*mRuby3*, Spec$^{R}$ | This study |
| AM004 | *vpvC*$^{W240R}$Δ*rbmA*Δ*bap1*Δ*rbmC*Δ*VC1807*::$P_{tac}$-*mNeonGreen*, Spec$^{R}$ | This study |
| AM007 | *vpvC*$^{W240R}$Δ*rbmA*Δ*VC1807*::$P_{tac}$-*SCFP3A*, Spec$^{R}$ | This study |
| AM008 | *vpvC*$^{W240R}$Δ*rbmA*Δ*bap1*Δ*rbmC*Δ*VC1807*::$P_{tac}$-*SCFP3A*, Spec$^{R}$ | This study |
| AM010 | *vpvC*$^{W240R}$Δ*rbmA*Δ*bap1*Δ*rbmC*Δ*VC1807*::$P_{tac}$-*SCFP3A*, Spec$^{R}$, pCMW121 | This study |
| AM011 | *vpvC*$^{W240R}$Δ*rbmA*Δ*bap1*Δ*rbmC*Δ*VC1807*::$P_{tac}$-*SCFP3A*, Spec$^{R}$, pCMW126 | This study |
| AM030 | *vpvC*$^{W240R}$Δ*rbmA*Δ*bap1*Δ*rbmC*Δ*VC1807*::$P_{tac}$-*mRuby3*, Spec$^{R}$ | This study |
| AM070 | *vpvC*$^{W240R}$Δ*rbmA*Δ*VC1807*::$P_{tac}$-*SCFP3A*, Spec$^{R}$, pJY26 | This study |
| AM082 | *vpvC*$^{W240R}$Δ*rbmA*Δ*rbmB*::Kan$^{R}$Δ*VC1807*::*SCFP3A*, Spec$^{R}$ | This study |
| AM041 | Δ*rbmA*Δ*bap1*Δ*rbmC*Δ*VC1807*::$P_{tac}$-*mNeonGreen*, Spec$^{R}$ | This study |
| AM042 | Δ*vpsL*Δ*VC1807*::$P_{tac}$-*mNeonGreen*, Spec$^{R}$ | This study |
| WN7666 | *vpvC*$^{W240R}$, BH625 | This study |
| WN7668 | *vpvC*$^{W240R}$Δ*rbmA*, BH625 | This study |
| **Plasmid** | | |
| pCMW121 | Kan$^{R}$, $P_{tac}$-*VC1086* (pEVS141 backbone) | [11] |
| pCMW126 | Kan$^{R}$, $P_{tac}$-*VC1086** (active-site mutant, pEVS141 backbone) | [11] |
| pJY26 | Kan$^{R}$, $P_{BAD}$-*rbmB* (pYS249 backbone) | This study |
| BH625 | Cm$^{R}$, $P_{vpsL}$-*luxCDABE* (pBBRlux backbone) | [12] |

**Table S2:** General coarse-grained parameters for bacterial cell model with their corresponding physical values.

| **Parameters** | **Simulation (reduced unit)** | **Physical** |
|---|---|---|
| Energy scale ($k_B T$) | $6.57 \times 10^{-4}$ | $4.14 \times 10^{-21}$ N m |
| Cell length | 30 | $3 \times 10^{-6}$ m |
| Cell diameter | 10 | $1 \times 10^{-6}$ m |
| Cell bead diameter, $\sigma_{cell}$ | 1.0 | 100 nm |
| Polymer bead diameter, $\sigma_{pb}$ | 1.0 | 100 nm |
| Cell shear modulus | 1.0 | $6.3 \times 10^{-2}$ N m$^{-1}$ |
| Cell area constant ($k_a$) | 0.075 | $4.72 \times 10^{-3}$ N m$^{-1}$ |
| Cell local area constant ($k_d$) | 3.67 | $2.31 \times 10^{-1}$ N m$^{-1}$ |
| Cell volume constant ($k_v$) | 0.3952 | $2490 \times 10^{2}$ N m$^{-2}$ |
| Cell bending constant ($k_b$) | 0.7937 | $5 \times 10^{-16}$ N m |
| Cut-off distance of LJ potential for pure repulsion | 1.1225 | 112.25 nm |
| Cut-off distance of LJ potential for attraction | 2.5 | 250 nm |
| Depth of the potential well of LJ potential for pure repulsion | $6.57 \times 10^{-4}$ | $4.14 \times 10^{-21}$ N m |
| Depth of the potential well of LJ potential induced by VPS production | $6.57 \times 10^{-5}$ | $4.14 \times 10^{-22}$ N m |
| Depth of the potential well of LJ potential for attraction | $3.28 \times 10^{-2}$ | $20.7 \times 10^{-20}$ N m |
| Temperature | 0.01 | 300 K |
| Timestep | 0.01 | 11.4 ns |
| Maximum run time | 100,000,000 | 1.14 s |

**Table S3:** Definitions of parameters, their notations and units used in the mathematical model.

| Notation | Parameter definition | Unit |
|---|---|---|
| $r_{\mathrm{P}}$ | Division rate of P cell | $\mathrm{hrs}^{-1}$ |
| $r_{\mathrm{nP}}$ | Division rate of nP cell | $\mathrm{hrs}^{-1}$ |
| $s$ | Relative advantage from being nP cell | 1 |
| $d_{\mathrm{P}}$, $d_{\mathrm{nP}}$ | Removal rates for P and nP cells | $\mathrm{hrs}^{-1}$ |
| $d$ | Basic death rates for both P and nP cells | $\mathrm{hrs}^{-1}$ |
| $d_{\mathrm{f}}$ | Wash-out rate due to flow, no sticking | $\mathrm{hrs}^{-1}$ |
| $A$ | Reduction in wash-out rate for a single P cell | 1 |
| $\upsilon$ | The normalization coefficient in the model of adhesion | 1 |
| $\mu_{\mathrm{P}\to\mathrm{nP}}$ | Conversion rate from P to nP cell | $\mathrm{hrs}^{-1}$ |
| $\mu_{\mathrm{nP}\to\mathrm{P}}$ | Conversion rate from nP to P cell | $\mathrm{hrs}^{-1}$ |
| $\mu$ | Max conversion rate from P to nP and from nP to P cell | $\mathrm{hrs}^{-1}$ |
| $\beta$ | Exponents in the expressions for the conversion rates | 1 |
| $N$ | Linear grid size | 2 μm |

**Supplementary References:**

1. Ye, H. *et al.* OpenFSI: A highly efficient and portable fluid–structure simulation package based on immersed-boundary method. *Comput. Phys. Commun.* **256**, 107463 (2020).
2. Plimpton, S. Fast parallel algorithms for short-range molecular dynamics. *J. Comp. Phys.* **117**, 1–19 (1995).
3. Zhang, Q. *et al.* Mechanical resilience of biofilms toward environmental perturbations mediated by extracellular matrix. *Adv. Funct. Mater.* **32**, 2110699 (2022).
4. Humphrey, W., Dalke, A. & Schulten, K. VMD: Visual molecular dynamics. *J. Mol. Graph.* **14**, 33–38 (1996).
5. Yan, J., Nadell, C. D. & Bassler, B. L. Environmental fluctuation governs selection for plasticity in biofilm production. *ISME J.* **11**, 1569–1577 (2017).
6. Gillespie, D. T. Stochastic simulation of chemical kinetics. *Annu. Rev. Phys. Chem.* **58**, 35–55 (2007).
7. Gibson, M. A. & Bruck, J. Efficient exact stochastic simulation of chemical systems with many species and many channels. *J. Phys. Chem. A* **104**, 1876–1889 (2000).
8. Secor, P. R., Michaels, L. A., Ratjen, A., Jennings, L. K. & Singh, P. K. Entropically driven aggregation of bacteria by host polymers promotes antibiotic tolerance in *Pseudomonas aeruginosa*. *Proc. Natl. Acad. Sci. USA* **115**, 10780 (2018).
9. Fernandez, N. L. *et al. Vibrio cholerae* adapts to sessile and motile lifestyles by cyclic di-GMP regulation of cell shape. *Proc. Natl. Acad. Sci. USA* **117**, 29046–29054 (2020).
10. Dalia, T. N., Chlebek, J. L. & Dalia, A. B. A modular chromosomally integrated toolkit for ectopic gene expression in *Vibrio cholerae*. *Sci. Rep.* **10**, 15398 (2020).
11. Waters, C. M., Lu, W., Rabinowitz, J. D. & Bassler, B. L. Quorum sensing controls biofilm formation in *Vibrio cholerae* through modulation of cyclic di-GMP levels and repression of *vpsT*. *J. Bacteriol.* **190**, 2527–2536 (2008).
12. Lenz, D. H., Miller, M. B., Zhu, J., Kulkarni, R. V. & Bassler, B. L. CsrA and three redundant small RNAs regulate quorum sensing in *Vibrio cholerae*. *Mol. Microbiol.* **58**, 1186–1202 (2005).
